# Using MAP-Elites to Optimize Self-Assembling Behaviors in a Swarm of Bio-micro-robots

**DOI:** 10.1101/845594

**Authors:** Leo Cazenille, Nicolas Bredeche, Nathanael Aubert-Kato

**Affiliations:** Department of Information Sciences, Ochanomizu University, Tokyo, Japan; Sorbonne Université, CNRS, Institut des Systèmes Intelligents et de Robotique, ISIR, F-75005 Paris, France

**Keywords:** Bio-micro-robots, swarm robotics, molecular programming, evolutionary robotics, quality-diversity algorithms, MAP-Elites

## Abstract

We are interested in programming a swarm of molecular robots that can perform self-assembly to form specific shapes at a specific location. Programming such robot swarms is challenging for two reasons. First, the goal is to optimize both the parameters and the structure of chemical reaction networks. Second, the search space is both high-dimensional and deceptive. In this paper, we show that MAP-Elites, an algorithm that searches for both high-performing and diverse solutions, outperforms previous state-of-the-art optimization methods.

## 1. INTRODUCTION

In this paper, we consider the challenge of automatically programming a swarm of micro-robots (≫ 1000 robots). The micro-robots we consider are microscopic agarose beads functionalized with bio-molecules (single strand DNA of 12-24 base pairs) [3]. The beads forming the body of the robots are small enough to move through Brownian motion. The DNA functionalization allows them to produce (a) chemical signals that may impact the behavior of nearby beads and (b) an anchoring signal that will attach them to their neighbors. Clusters created from aggregated beads move slower or even stop, based on their size. That property allows us to control where the beads should go, which we use to create self-assembled structures. Thanks to that scale and the low price and availability of the molecular components, we previously designed and validated *in vitro* a simple system of a million micro-robots that self-assemble [1].

We previously introduced the BioNeat algorithm [1], inspired from state-of-the-art Neat algorithm [7]. BioNeat aims at producing chemical reaction networks (CRN) which represent bio-molecules to be attached to beads or left floating in the 2-dimensional substrate. Given a target for self-assembly (*e.g.*, in Fig. 1), the objective is that micro-robots assemble to one another in the target area.

**Fig. 1.**
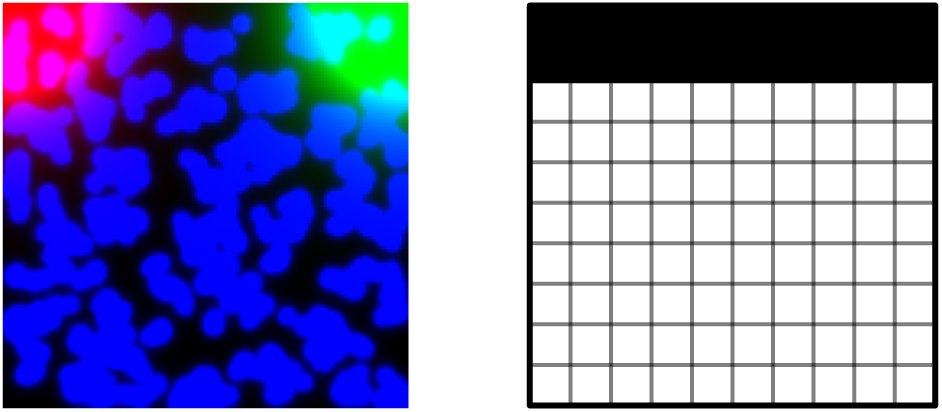
**Left**: Simulation of an experiment where a swarm of micro-beads (in blue) interact with their environment and self-aggregate. Two gradient sources, in the top-left and top-right positions, are continuously diffused isotropically in the environment. They are shown respectively in red and green. **Right**: Target for self-assembly. Beads should aggregate in the black area.

Though initial results are promising, this previous work revealed an important limit of the BioNeat algorithm. Due to the complex search space, the algorithm quickly converged to sub-optimal results even when relatively simple assembled shape where considered (e.g. self-assembling into a single region placed in the center of the environment).

In order to tackle this, we introduce the use of MAP-Elites [5], an illumination algorithm that favors exploration over pure optimization, thus reducing the risk of premature convergence during the search. We also implement a new mechanism, termed *topological initialization* that helps MAP-Elites to bootstrap exploration when confronted with search spaces with large neutral regions.

The following, we describe the Methods, both in terms of experimental setup, objective function and optimization algorithms. Then, we present the results comparing the original BioNeat algorithm with MAP-Elites, and show that the latter dominates both in quality and speed of convergence.

## 2. METHODS

We consider a swarm comprised of molecular robots made from spherical beads grafted with specific strands of DNA. Those strands are of two types: (a) templates used to capture signal DNA from the environment, process it and produce signal back; (b) anchoring points, which are used for aggregation when the anchoring signal is present. In this Section, we summarize the general idea behind the model. A detailed description, including a validation with *in vitro* experiments, can be found in our previous work [1].

### 2.1. Model

Processing and production of signals are based on the PEN DNA toolbox [4]. Using this toolbox, signal strand can attach to a compatible template and either triggers the production of a different DNA strand (activation) or prevents activation from other strands (inhibition). Signal strands are spread in the environment through chemical diffusion and are degraded over time through enzymatic activity.

Robots move through Brownian motion: individual robots move much faster than aggregated clusters, which in turn may become completely immobilized when they get large enough.

Beads are modelled as disks moving in the environment through Brownian motion. While the simulation is 2-dimensional, we consider that the beads are moving in a 3-dimensional environment, which allows us to ignore collisions. In presence of anchoring signal, beads have a probability to aggregate. That probability is simulated by implementing a Gillespie-like step: considering the current concentration, it is possible to predict when the next aggregation event will occur. If that event happens before the next time step, aggregation is considered successful. The reverse reaction, having a bead separating from an aggregate, is computed the same way.

The local concentration of signal-producing species is directly proportional to the number of beads at a given point of space. That is, we sum the concentrations of DNA molecules grafted on all the beads that are present. Note that, due to the non-linearity of the enzymatic reactions involved, a linear increase in signal-producing species does not necessarily mean a linear increase in the production of signal species. Finally, the production, diffusion, and degradation of signal species are modeled through reaction-diffusion. For a given signal *S*, we have:

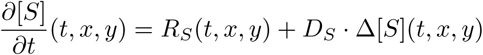

where [*S*] is the concentration of *S*, *R*_*S*_ is the contribution from reactions (*e.g.*, production or degradation), *D*_*S*_ is the diffusion coefficient of *S* and ∆ is the Laplacian operator.

We define a target for self-assembly corresponding to the black area shown in Fig. 1. It is composed of a single line at the top position of the arena. Our objective is for the micro-beads to self-assemble into the target area, shown in black, starting from their initial random positions. Two fixed gradient sources are arranged in the top-left and top-right positions in the arena (Fig. 1 left). They each emit a respective type of signal strand that diffuse throughout the environment. They provide the micro-beads information about their localization, potentially inducing self-assembly. As the gradients diffuse isotropically, self-assembling into straight patterns can be difficult.

We quantify the performance of a simulation by using the method described in [1]. The experimental arena is discretized into a *N × N* matrix of cells, with *N* = 160. This allows to compute the following *match-nomatch* score:

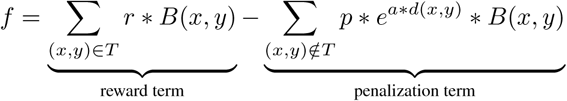

with *T* the set of (*x, y*) positions in the target area, *r* a reward parameter, *B*(*x, y*) a Boolean function of the presence of micro-beads at position (*x, y*), *d* the distance of a cell towards the closest position in *T* and *a* a scaling parameter. Individuals are rewarded according to the number of cells within *T* and penalized for cells outside of *T*, with penalization increasing with distance to the target area. Table 1 lists the parameters used for simulation and fitness computation. In order to take into account noise during evaluation, each candidate solution is reevaluated 5 times to provide a reliable estimation of its performance.

Individuals are deemed valid only if their respective topology matches the requirements of Table 2 in term of number of nodes and number of activation and inhibition templates.

**Table 1.** Simulations and fitness parameters.

| Simulation Parameter | Value |
| --- | --- |
| Arena size | 1mm × 1mm |
| Beads | 500 |
| Bead size (aggregation) | 50μm |
| Bead size (Brownian motion) | 5μm |
| Temperature | 43°C |
| Grid size | 160 × 160 |
| Time discretization | 0.1 min per step |
| Simulation duration | 1000 steps (*i.e.*, 100 min) |
| Target width | 0.20mm (20% of arena width) |
| Fitness Parameter | Value |
| $r$ (reward) | 1.0 |
| $p$ (penalty) | 0.2 |
| $a$ (scaling) | 0.1 |

**Table 2.** Parameters of BioNeat and MAP-Elites.

| Optimization Parameter | Value |
| --- | --- |
| Evaluation Budget | 3000 |
| Number of retrials per individuals | 5 |
| Maximal number of activation nodes | 6 |
| Maximal number of activation templates | 7 |
| Maximal number of inhibition templates | 7 |
| BIONEAT Parameter | Value |
| Target number of BIONEAT species | 20 |
| Population Size | 50 |
| Number of templates | 1 – 13 |
| MAP-Elites Parameter | Value |
| Shape of the grid | 7 |
| Batch size | 50 |
| Number of elites per grid bin | 1 |
| Number of templates | 7 – 13 |
| Mutation operator | Probability |
| Parameter mutation | 0.80 |
| Add template strand | 0.05 |
| Remove template strand | 0.05 |
| Add signal species | 0.05 |
| Add inhibition species | 0.05 |

### 2.2. Optimization

We rely on two methods to optimize the structure of chemical reaction networks: BioNeat and Map-Elites. BioNeat is an evolutionary algorithm we first introduced in [1]. It takes inspiration from the famous state-of-the-art Neat algorithm [7], which was originally designed to optimize artificial neural networks. BioNeat uses specific variation operators to navigate the search space of chemical reaction networks. It is also capable to protect innovation, that is to explore several regions of the search space simultaneously, balancing between novelty of a particular design and the quality of solutions. We previously showed that BioNeat provides efficient solutions for targets comprised of a single regions (horizontal or vertical lines, see [1] for details). The main limit of BioNeat is that while it can protect innovation for some time, there is not guarantee that it is capable of escaping completely the curse of premature convergence due to the pressure for optimizing towards (possibly only temporary) better solutions.

BioNeat searches through CRN topologies iteratively. This behavior is inherited from Neat that postulated that iterated small changes in topologies would often only result in a mild effect on the fitness values. It may not be the case with CRN, where small changes in topology can have severe effects in fitness values, a particular trait of deceptive and hard-explore problems. This may explain why BioNeat is prone to premature convergence during optimization, as it does not possess a way to explore a totally unexplored niche in the space of topologies. While this aspect is partially mitigated with BioNeat mechanism of speciation, which allows the parallel optimization of several topological niches, it does not enforce the discovery of totally novel niches. Worst, as new species are only created through atomic mutations (*i.e.*, change only a small part of the topology), the surviving species may contains individuals with very similar topologies, possibly in the same topological niche.

In order to favor exploration over pure optimization, we rely on MAP-Elites [5], which is an illumination algorithm (also sometimes referred to as Quality-Diversity algorithm [6]) that decomposes the search space into regions based on feature descriptions. It considers how does a candidate solution *look like* in the phenotypic space instead of considering how it is coded in the genotypic space. This method is particularly suited to cope with multi-modal, deceptive, hard-explore and ill-defined problems where traditional optimization algorithms would be prone to premature convergence, as in evolutionary robotics [5]

MAP-Elites iteratively regroups the explored solutions in a grid of elites. This results in a collection of high-performing individuals across a number of features selected by the user, corresponding to the axes of the grid. Here, we only consider a single feature corresponding to the total number of templates in the topology of an individual. CRN with smaller number of templates are easier to test experimentally but lose expressivity. Conversely, large-sized CRN can describe more complex behaviors, which may be necessary for the beads to successfully self-aggregate into the target shape. As such, a trade-off in term of topology complexity has to be considered, possibly after the optimization process. This substantiates methodologies that concurrently search for topologies of differing sizes.

MAP-Elites is equipped with the same set of mutation operators as BioNeat, and also retains its capability to optimize iteratively the topologies. We describe a novel methodology to bootstrap MAP-Elites exploration by initializing a collection of individuals with random topologies. This approach allows MAP-Elites to consider a large number of differing topologies from the start. It makes use of the BioNeat mutation operators to generate individuals of varying topologies across a range of number of templates. In one optimization run, 10% of the individuals are initialized with a random topology (*i.e.*, 300 individuals for an evaluation budget of 3000). This contrasts with the BioNeat approach, where only small iterative changes in topologies are possible from mutations, and where individuals with totally new topologies are not initialized.

Table 2 lists the chosen parameters for the MAP-Elites algorithm. We use our own implementation of BioNeat and MAP-Elites. BioNeat is open source and available from https://bitbucket.org/AubertKato/bioneat/. MAP-Elites is coded in Python using the QDpy library [2] and freely available as open source software at https://gitlab.com/leo.cazenille/qdpy. All additional scripts used in this paper are available at https://bitbucket.org/leo-cazenille/daccad-qd.

## 3. RESULTS

The BioNeat and MAP-Elites algorithms are used to optimize CRNs on the target described earlier, with a budget of 3000 evaluations per run, and with 16 replicates.

For each setting, the evolution of fitness values during 16 optimization runs are presented in Fig. 2. As shown in the Figure, MAP-Elites dominates BioNeat in both performance and speed of convergence during the course of optimization. This is confirmed by an additional experiment, comparing methods using the best individuals from each run of each method (16 individuals per method). We used a Mann-Whitney U test on the distribution of fitness of their respective best individuals. Results are summarized in Table 3. Again, MAP-Elites dominates BioNeat, with a *p* value of 10 − 6 (Mann-Whitney U Test).

**Table 3.** Fitness scores of the best-performing individuals for BioNeat and MAP-Elites, across 100 retrials.

|  |  |
| --- | --- |
| BIONEAT | $0.71 \pm 0.03$ |
| MAP-Elites | $0.87 \pm 0.03$ |

**Fig. 2.**
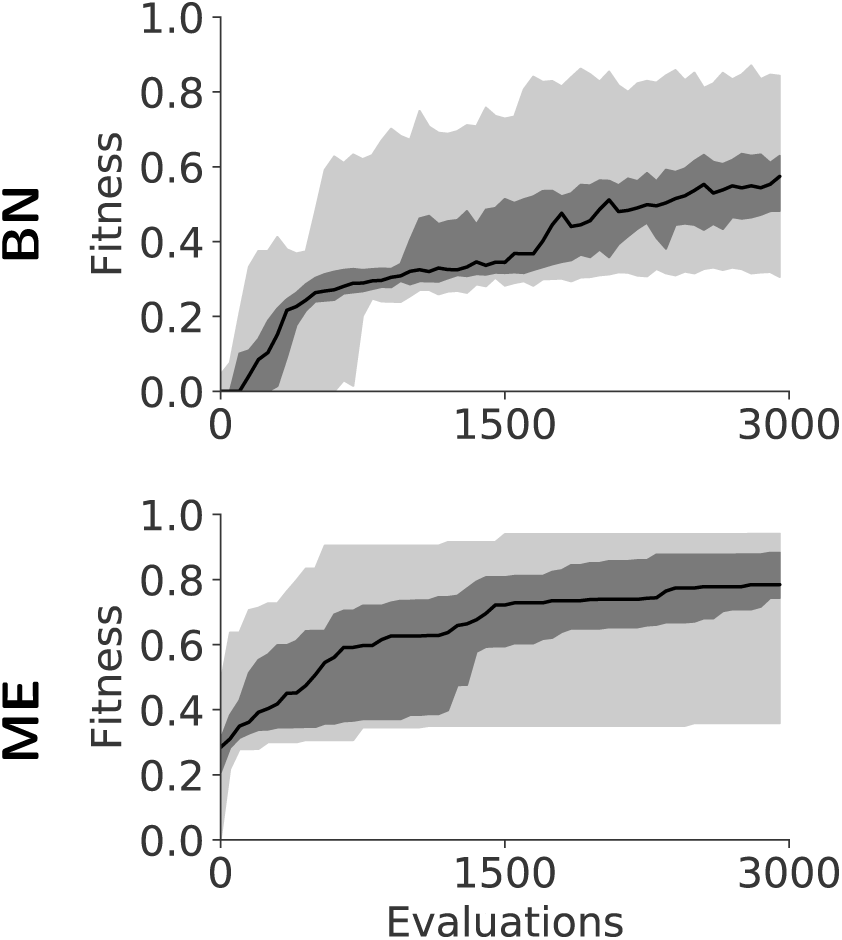
Evolution of the median quality of the best-performing individuals for each method (BN: BioNeat; ME: MAP-Elites). Optimization methods are tested across 24 different runs. A fitness of 1.0 corresponds to the best performance. The darker shade represents the 25 to 75 percentiles. The lighter shade encompasses the minimal and maximal values.

Figure 3 shows the final state of a full simulation of the best-performing solutions for each optimization method. For clarity, only the anchoring signal is represented, corresponding to large self-assembled clusters of micro-beads.

**Fig. 3.**
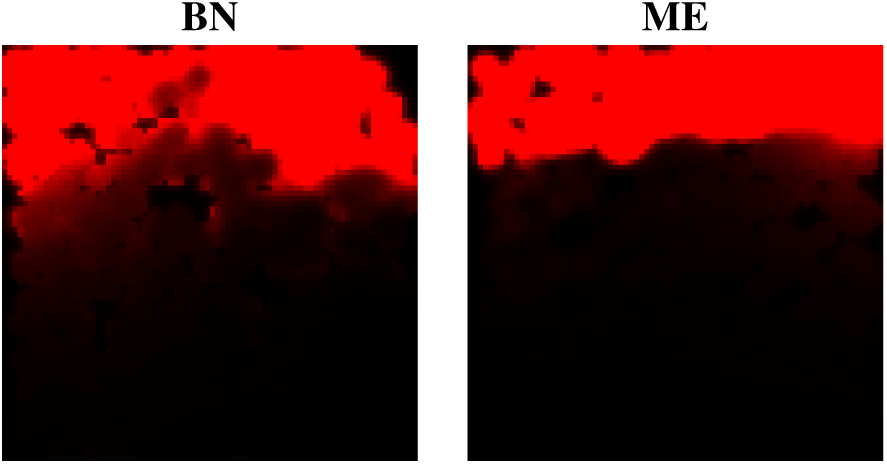
Examples of final states obtained by the best-performing individuals optimized by both optimization methods

## 4. DISCUSSION AND CONCLUSION

We introduced the MAP-Elites algorithm to optimize chemical reaction networks of a swarm of bio-micro-robots to self-assemble into a target shaped target. While MAP-Elites is originally a quality-diversity algorithm that emphasize exploration over exploitation, we showed that the incentive for exploration can directly benefits optimization in a setting involving complex interaction between micro-robots.

As short term perspectives, we are investigating the benefits of using MAP-Elites for more complex target shapes. In particular, we propose to refine MAP-Elites’ feature descriptors to capture useful phenotypical traits at the level of the robot swarm, so as to guide exploration towards relevant behavioural patterns.

Secondly, the current simulator we use is highly reliable with respect to simulating real chemical reactions, but results in a high computational cost. We aim to tackle this problem by introducing a surrogate model to trade speed over accuracy. As such approximation cannot capture the details of real chemistry, we propose to mix the use of surrogate model and realistic simulation to speed up optimization while retaining high quality solution.

## ACKNOWLEDGEMENTS

This work was supported by JSPS KAKENHI Grant Number JP17K00399 and by Grant-in-Aid for JSPS Fellows JP19F19722. This work was also supported by the Agence Nationale pour la Recherche under Grant ANR-18-CE33-0006 (MSR project).

